# Systems genetic discovery of host-microbiome interactions reveals mechanisms of microbial involvement in disease

**DOI:** 10.1101/349605

**Authors:** Jason A. Bubier, Vivek M. Philip, Christopher Quince, James Campbell, Yanjiao Zhou, Tatiana Vishnivetskaya, Suman Duvvuru, Rachel Hageman Blair, Juliet Ndukum, Kevin D. Donohue, Charles Phillips, Carmen M. Foster, David J. Mellert, George Weinstock, Cymbeline T. Culiat, Erich J. Baker, Michael A. Langston, Bruce O’Hara, Anthony V. Palumbo, Mircea Podar, Elissa J. Chesler

## Abstract

The role of the microbiome in health and disease involves complex networks of host genetics, genomics, microbes and environment. Identifying the mechanisms of these interactions has remained challenging. Systems genetics in the laboratory mouse enables data-driven discovery of network components and mechanisms of host-microbial interactions underlying multiple disease phenotypes. To examine the interplay among the whole host genome, transcriptome and microbiome, we mapped quantitative trait loci and correlated the abundance of cecal mRNA, luminal microflora, physiology and behavior in incipient strains of the highly diverse Collaborative Cross mouse population. The relationships that are extracted can be tested experimentally to ascribe causality among host and microbe in behavior and physiology, providing insight into disease. Application of this strategy in the Collaborative Cross population revealed experimentally validated mechanisms of microbial involvement in models of autism, inflammatory bowel disease and sleep disorder.

**eTOC Blurb:** Host genetic diversity provides a variable selection environment and physiological context for microbiota and their interaction with host physiology. Using a highly diverse mouse population Bubier et al. identified a variety of host, microbe and potentially disease interactions.

**Highlights:**

* **18 significant species-specific QTL regulating microbial abundance were identified**
* **Cis and trans eQTL for 1,600 cecal transcripts were mapped in the Collaborative Cross**
* **Sleep phenotypes were highly correlated with the abundance of *B.P. Odoribacter***
* **Elimination of sleep-associated microbes restored normal sleep patterns in mice.**

## Introduction

Although the human microbiome has been implicated as an important factor in health and disease (Wen et al., 2008), the mechanisms by which it influences human physiology are largely unknown. Experiments that manipulate specific genetic, molecular and microbial components of the microbe-host interface are essential to dissection of these mechanisms (Vijay-Kumar et al., 2010), but identification of targets for experimental manipulation remains a significant challenge. However, both microbial community composition and its effects on host health are modulated by host characteristics that exhibit heritable variation (Benson et al., 2010; Campbell et al., 2012; McKnite et al., 2012; Snijders et al., 2016), providing the opportunity to identify genetic variants and associated traits to serve as entry points for investigating key functional pathways at the microbe-host interface (Knights et al., 2014; McKnite et al., 2012; Willing et al., 2010).

Studies of the gut microbiome have produced convincing evidence for a microbial influence over many host traits including human gastrointestinal (GI) disorders (Knights et al., 2014; Machiels et al., 2014; Willing et al., 2010), metabolic traits, diabetes (Vijay-Kumar et al., 2010; Wen et al., 2008) and obesity (Carlisle et al., 2013; McKnite et al., 2012; Parks et al., 2013). Perhaps more surprising is the influence of gut microbiota and the metabolites they produce on brain and behavior (Bravo et al., 2011; Carter, 2013; Lewin et al., 2011). Despite the importance of these microbial influences, the mechanisms of these interactions often remain unknown.

There are many well-documented relationships among host genetic variation, intestinal flora composition and disease reported in human genetic analyses (Deloris Alexander et al., 2006; Goodrich et al., 2014; Jacobs and Braun, 2014; Knights et al., 2014; Turnbaugh et al., 2009). Because mice and humans harbor similar microbiota at high taxonomic levels (Krych et al., 2013; Ley et al., 2008), systems genetic analysis in laboratory mice can be an effective tool for discovering the mechanisms of host−microbe interaction in a large-scale, data driven manner. This quantitative genetic approach provides a means of holistic assessment of the relations among host, microbe and disease through the use of population genetic variation, one of the greatest determinants of microbial community composition in mice (Campbell et al., 2012; Deloris Alexander et al., 2006). The study of natural genetic variation (Campbell et al., 2012) and engineered mutations (Spor et al., 2011; Turnbaugh et al., 2006), also enable deep dissection of the biology of the microbiome and discovery of host genetic loci that regulate microbial abundance (Benson et al., 2010; McKnite et al., 2012). The transcriptome provides insight into the host microenvironment by quantifying the relative abundance of transcripts encoding host pathways involved in metabolic responses, the production and presentation of cell-surface antigens, and constituents of the immune system such as the gut associated lymphoid tissue, among other host processes that both shape and respond to gut microbiota.

The Collaborative Cross (CC) mouse population, (Chesler et al., 2008; Churchill et al., 2004) constructed from the cross of eight diverse inbred progenitor strains, was designed for high precision, (Philip et al., 2011) and high diversity systems genetic analysis. The host genetic variation among this population results in diverse microbiome compositions (Campbell et al., 2012) and physiological and behavioral phenotypes. Genetic correlations among these characteristics are used to construct systems genetic networks (Figure 1). Interrogation of these networks at the level of transcript, microbe and phenotypes enable the study of mechanisms of microbiota influence on health and disease by identifying causal mechanisms responsible for phenotypic correlations.

**Figure 1.**
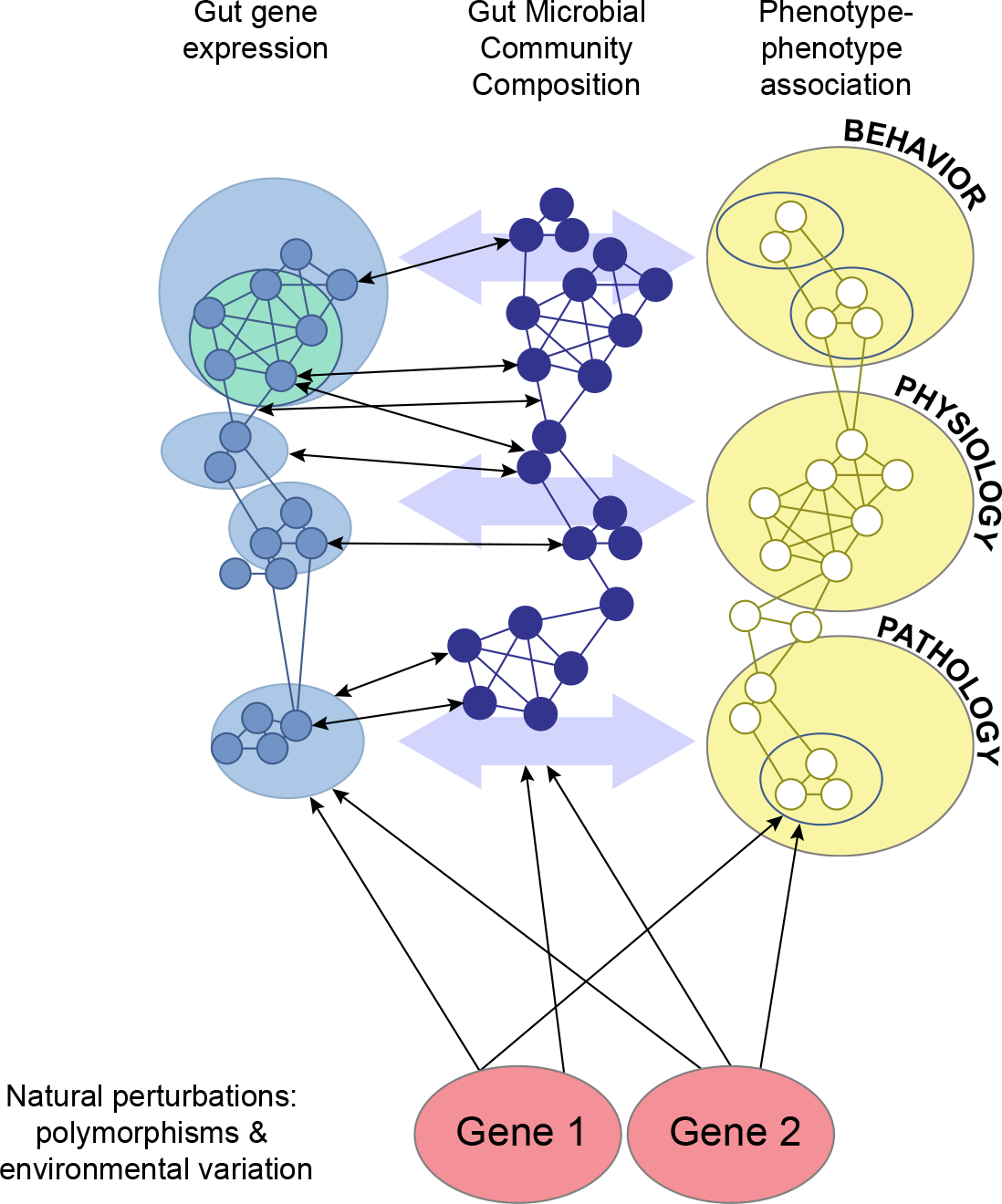
The system genetic model and gut microbiomics. Genetic variation in a host population together with the environment interact to affect gene expression in the host, microbial abundance and disease related traits. There is significant bi-directional interplay among the microbiome, host gene expression and disease related phenotypes, however, the effect of host genotype is unidirectional, and therefore causal.

Here, we integrated data for host genotype, disease related phenotypes, gut−microbiome composition and associated gut gene expression to develop a systems genetic network for the gut-microbiome interaction and its effects on host health. Specifically, we performed an integrative analysis leveraging cecal mRNA levels, luminal microbiome, physiology and behavior from over 100 incipient strains of the CC population. Through the analysis of relations among these measurements, we apply systems genetic analysis to problems in autism-related behaviors, inflammatory bowel disease and sleep.

## Results

### Microbial community composition of incipient CC Mice

We first determined the cecal microbial community composition of 206 CC mice of both sexes and 102 breeding lines using 454 pyrosequencing of amplicon libraries of the V4 region of the 16S SSU (Small Subunit) rRNA gene, revealing 13,632 Operational Taxonomic Units (OTUs). Taxonomic analysis of all sequences using the Ribosomal Database Project naïve Bayesian rRNA classifier (Cole et al., 2009; Cole et al., 2014) indicated a bacterial diversity similar to previously observed communities (Campbell et al., 2012)—Firmicutes comprised 89% of the microbial community and Bacteroidetes (9%) were the second most abundant phylum (Figure S1). In our previous study of replicate mice from the eight CC progenitor strains, we detected more phyla in the founders, but we show here that there are similar predominating phyla (Campbell et al., 2012) in the CC. The median broad-sense heritability of microbial abundance estimated by intraclass correlation in the CC founder strains for each OTU (Table S1) was 0.170, with 339 OTUs having a H^2^ > 0.3, indicating sufficiently heritable abundance for genetic mapping.

### Microbial Abundance QTLs

We performed quantitative trait locus (QTL) mapping to identify host genetic loci accounting for heritable variation in microbial abundance. There were eighteen statistically significant (q <0.05) microbial abundance (*Micab*) QTL (Table 1) among the mapped microbial OTU abundances. The 1.5 logarithm of odds (LOD) confidence intervals for the significant QTL range from 2 to 24 Mb in size, with an average size of 7.5 Mbp. The size is consistent with previous mapping studies in the CC breeding population (Philip et al., 2011; Snijders et al., 2016) and substantially smaller than conventional experimental crosses. This interval size coupled with the extensive genomic data becoming available for the CC founder population enables refinement of the QTLs down to the level of genes and variants in some cases.

**Table 1.** Significant QTLs for microbial abundance in the cecum of incipient CC mice.

| **MGI**<br>**QTL**<br>**Symbol** | **QTL Name** | **Chr** | **Peak**<br>**LOD**<br>**Score** | **Peak**<br>**Marker** | **Position**<br>**Mm 9 (bp)** | **1.5 LOD Interval** |  | **p-value** | **Size**<br>**(Mbp)** | **Gene**<br>**Weaver**<br>**GSID** |
| --- | --- | --- | --- | --- | --- | --- | --- | --- | --- | --- |
| Micab1 | Microbial abundance of Clostridiales<br>Ruminococcaceae Oscillibacter 1 | 1 | 8.20 | rs32084678 | 13,279,810 | rs6275656 | rs31653681 | 0.0033 | 6.34 | 217070 |
| Micab2 | Microbial abundance of Bacteroidales<br>Porphyromonadaceae Paludibacter 2 | 3 | 8.23 | rs31103355 | 108,854,325 | rs31431100 | rs37044521 | 0.0029 | 4.09 | 217071 |
| Micab3 | Microbial abundance of Clostridiales<br>Lachnospiraceae Marvinbryantia 3 | 3 | 8.13 | rs30089246 | 37,569,141 | rs30552223 | rs30158956 | 0.0037 | 2.03 | 217072 |
| Micab4 | Microbial abundance of Clostridiales<br>Lachnospiraceae Roseburia 4 | 4 | 9.57 | rs32690134 | 136,028,098 | rs27619452 | rs3685172 | 0.0005 | 9.45 | 217077 |
| Micab5 | Microbial abundance of<br>Coriobacteriales Coriobacteriaceae<br>Enterorhabdus 5 | 5 | 8.84 | rs6377391 | 119,128,609 | rs29633871 | rs6354701 | 0.0009 | 4.65 | 217078 |
| Micab6 | Microbial abundance of Clostridiales<br>Lachnospiraceae Sporobacterium 6 | 5 | 8.42 | rs8265964 | 138,359,981 | rs32246505 | rs32318125 | 0.002 | 4.00 | 217079 |
| Micab7 | Microbial abundance of Bacteroidales<br>Porphyromonadaceae Odoribacter 7 | 7 | 8.84 | rs31494696 | 77,651,351 | rs33107817 | rs6373775 | 0.0009 | 3.53 | 217080 |
| Micab8 | Microbial abundance of Clostridiales<br>Ruminococcaceae Lactonifactor 8 | 7 | 8.88 | rs47611520 | 47,761,932 | rs3661776 | rs6176297 | 0.0009 | 13.55 | 217081 |
| Micab9 | Microbial abundance of Bacteroidales<br>Porphyromonadaceae Odoribacter 9 | 7 | 8.60 | rs31494696 | 77,651,351 | rs33107817 | rs6373775 | 0.0013 | 4.00 | 217082 |
| Micab10 | Microbial abundance of Clostridiales<br>Lachnospiraceae Anaerostipes 10 | 8 | 9.61 | rs32936112 | 47,123,375 | rs6281843 | rs31252778 | 0.0003 | 14.36 | 217083 |
| Micab11 | Microbial abundance of Clostridiales<br>Incertae Sedis XIV Blautia 11 | 8 | 9.49 | rs33429737 | 31,919,239 | rs6399870 | rs50110045 | 0.0005 | 5.19 | 217084 |
| Micab12 | Microbial abundance of Clostridiales<br>Clostridiaceae Caminicella 12 | 9 | 8.29 | rs30372085 | 80,440,479 | rs30432532 | rs33695839 | 0.0029 | 7.37 | 217092 |
| Micab13 | Microbial abundance of<br>Erysipelotrichales Erysipelotrichaceae<br>Turicibacter 13 | 10 | 8.87 | rs29327022 | 88,018,183 | rs6338556 | rs6265280 | 0.0009 | 24.62 | 217093 |
| Micab14 | Microbial abundance of Bacteroidales<br>Bacteroidaceae Bacteroides 14 | 11 | 8.14 | rs26971743 | 58,783,410 | rs6314621 | rs26972849 | 0.0036 | 6.30 | 217094 |
| Micab15 | Microbial abundance of Clostridiales<br>Lachnospiraceae Syntrophococcus 15 | 15 | 9.47 | rs6388530 | 93,634,974 | rs31931586 | rs49819430 | 0.0005 | 10.43 | 217095 |
| Micab16 | Microbial abundance of Bacteroidales<br>Porphyromonadaceae Tannerella 16 | 19 | 8.10 | rs30320578 | 47,732,625 | rs36280504 | rs30760881 | 0.004 | 6.39 | 217096 |
| Micab17 | Microbial abundance of Clostridiales<br>Ruminococcaceae<br>Hydrogenoanaerobacterium 17 | X | 8.50 | rs6292190 | 155,749,346 | rs29276152 | rs29306363 | 0.0017 | 5.59 | 217097 |
| Micab18 | Microbial abundance of Clostridiales<br>Lachnospiraceae Lachnobacterium 18 | X | 8.20 | rs6213950 | 163,790,061 | rs8255374 | rs31682358 | 0.0033 | 3.01 | 217098 |

### Expression QTLs (eQTL) in the CC Cecum

To characterize the host intestinal state, we profiled mouse mRNA abundance in the cecal tissue surrounding the microbial sample. Transcript abundance estimates were generated for 36,308 microarray probes, representing 27,149 genes. Heritability of transcript abundance exceeded H^2^=0.3 for 1,990 probes in the founder populations. QTL analysis was performed to identify host genomic regions harboring allelic variants that influence the abundance of each probe, resulting in detection of statistically significant QTL (q< 0.05) for 1,641 probes, corresponding to 1,513 genes (Table S2). Of these, 950 loci (57.9%) are cis-eQTL (Figure S2), which contain polymorphisms that are proximal to transcript coding regions. Such loci are useful in identifying expression regulatory mechanisms in the effect of genetic variation on complex traits.

### Genetic correlation analysis reveals relations among genes and microbes

The characterization of genetically heterogeneous populations enables the multi-scale correlation and clustering of high-dimensional phenotypic data across individuals, including gut microbial abundance for all detected OTUs, intestinal-expressed genes, and disease-related phenotypes.

For each of 393 microbial OTUs subject to genetic analysis, a set of co-abundant transcripts were detected across individuals (Table S3). Biclustering of gene-microbe correlations using a biclique enumeration algorithm (Zhang et al., 2014) revealed 318 clusters of multiple cecal microbes and the set of co-abundant transcripts found within the cecal intestine in each cluster. Sets of microbes and associated transcripts were deposited in the GeneWeaver database for access and subsequent comparative analysis with other heterogeneous functional genomic data in the form of gene lists.

These sets of correlated transcripts, microbes and disease measures were interrogated along with the microbial QTL and intestinal eQTL to identify the elements of biological networks of host and microbial interactions that were subject to experimental perturbation. From the interrogation of these relationships in systems genetic networks, we extracted specific mechanistic hypotheses for microbe-host interactions. In each of three examples that follow, data were interrogated by starting from a microbial QTL, microbe-transcript cluster, or microbe-physiology trait correlation, respectively, revealing putative relations that could begin to be tested experimentally.

### Genetic characterization of disease-associated microbes reveals potential mechanism of microbial influence of autism-related behavior

Microbial abundances, (*Micab*) QTLs, and cecal eQTLs were integrated to elucidate the mechanism underlying a previously reported association between a microbe, *Enterorhabdus* and autism spectrum disorder (ASD). Some microbial species, including *Enterorhabdus* are more abundant in the valproic acid (VPA)-induced mouse model of autism than non-exposed controls, a phenomenon associated with cecal butyrate production (de Theije et al., 2014). Levels of short-chain fatty acids such as butyric acid have also been associated with other autism models (Kratsman et al., 2016; Morris et al., 2016). Aberrant social behavior, a hallmark of ASD in humans, has been observed in germ free mice (Desbonnet et al., 2014). We found that *Enterorhabdus* is regulated by the *Micab5* locus. We hypothesized that genetic variants influencing abundance of *Enterorhabdus* may also influence ASD-related behaviors.

The microbial abundance QTL for *Enterorhabdus* (*Coriobacteriales Coriobacteriaceae*), *Micab5*, maps to Chromosome 5, an interval of 4.65 Mbp containing 121 genes (Figure 2A). Identification of cis-eQTLs, expressed genes regulated by local genetic variation, may help to identify the pathways and processes responsible for the QTL effect, and when there is phenotype and gene expression correlation a regulatory mechanism is most likely. To determine whether any of the genes in the interval are regulated by local expression regulatory polymorphisms, we intersected the list of cecal eQTLs with the 121 genes within the *Micab5* confidence interval, revealing two genes with significant cis-eQTL, *Tmem116* (probe34138 or ILMN_2624031) and *Trafdl* (probe_34454 or ILMN_2833441). In contrast to *Tmem116*, *Trafd1* is also a gene expression correlate of *Enterorhabdus* abundance (Spearman’s rho=0.1465, q = 0.0437) <u>and</u> the *Trafd1* eQTL allelic effect is similar to that of *Micab5* (Figure 2B-C). Therefore, *Trafd1* is a genetically plausible candidate. A search of Mammalian Phenotype Ontology (MP) annotations within Mouse Genome Database revealed that *Trafd1* interacts with bacterial and viral pattern recognition receptors (*e.g.*, toll-like receptors) thus *Trafd1* is a functionally plausible candidate gene for controlling microbial abundance. A search of Comparative Toxicogenomic Database (Davis et al., 2015; de Theije et al., 2014) revealed that VPA and acetaminophen are the top interacting partners of *Trafd1*, and that its expression is increased by VPA exposure (Jergil et al., 2011; Kultima et al., 2010). Thus, we hypothesize that perturbation of *Trafd1* abundance (through the cis regulatory QTL in the CC population) mimics the effects of VPA exposure on microbial abundance in the VPA autism model. In a test of this hypothesis, *Trafd1* knock-out mice were compared to controls on the three-chambered social interaction test, one test of ASD like behavior applied to the VPA model. There was a significant genotype effect such that knock-out mice spent less time interacting with a stranger mouse and more with the object than controls (F_genotype(1,27)_ = 7.9864, p=0.008). (Figure 2D). Strikingly, 16S rDNA sequencing of cecal contents of *Trafd1* knock-out and controls detected three OTUs corresponding to *Enterorhabdus*, the abundance of which did not differ across sexes or genotypes. However, between the genotype there were only four microbial taxa that differed in their abundance (Figure S3), one of which was *Butyrivibrio*, a known butyric acid-producing organism. One interpretation is that the QTL mapped for *Enterorhabdus* abundance may have been a surrogate for differential butyric acid production, because the *Butyrivibrio* abundance was decreased in the *Trafd1* knock-out mice but there was no difference in *Enterorhabdus*. Perhaps the allelic variation in *Trafd1* is associated with the abundance of microbes that can produce butyrate with downstream effects on social behavior, suggesting that *Trafd1* null mutants may represent a new model for ASD and warrant further study.

**Figure 2-.**
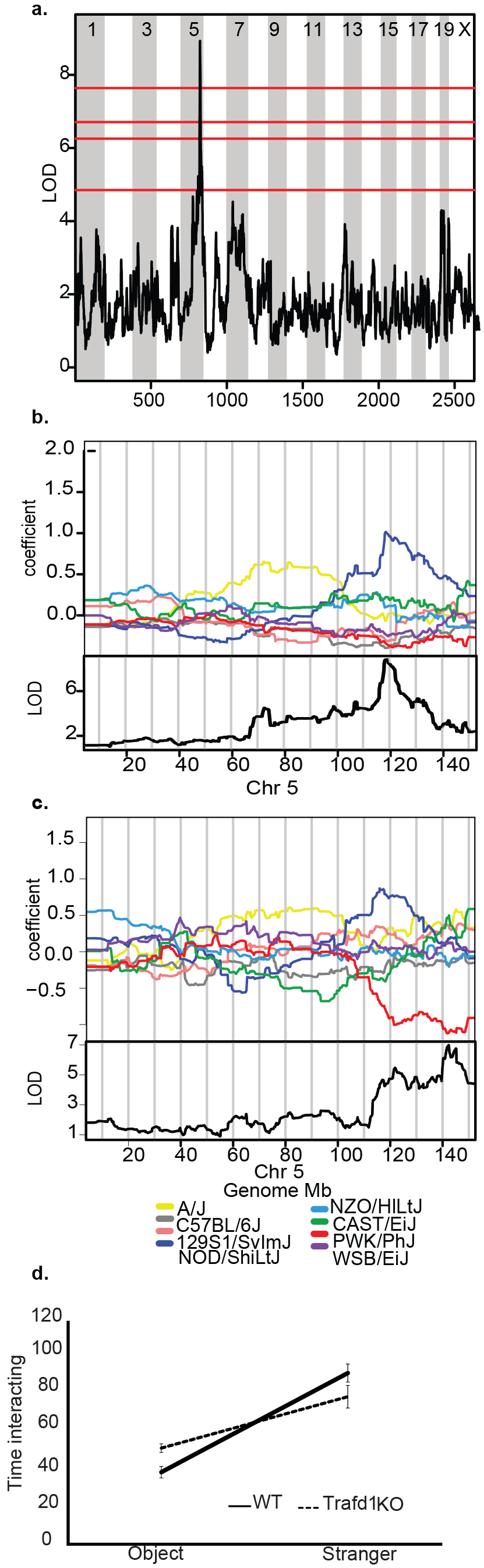
*Enterorhabdus* abundance in the cecum. a) Genome scan showing a large significant QTL (p<0.01) on chromosome 5 for regulation of *Enterorhabdus* abundance. Horizontal lines represent permutation based significance thresholds from the top down highly significant p<0.01, significant, p<0.05, highly suggestive, p<0.1, suggestive, p<0.63 b) Detailed map of Chromosome 5. Allelic effect plots of eight founder coefficients of the QTL mixed model representing the effect of each founder haplotype on phenotype. The NOD/LtJ allele on Chromosome 5 is associated with increased abundance c) Overlapping proximal (cis) eQTL for *Trafd1* expression on chromosome 5 with a similar NOD/LtJ driving increased expression. d) *Traf1*-KO mice spent more time interacting with the object, and less time with the stranger than their wild-type liter mates.

### Mapping microbes onto human disease through cross-species integrative functional genomics reveals host gene and microbial mechanisms associated with Inflammatory Bowel Disease

Sets of genes associated with particular OTUs were interrogated using GeneWeaver’s (Baker et al., 2012; Baker et al., 2009) tools for cross-species integrative functional genomics. In this approach, sets of genes associated with biological concepts such as disease state or molecular pathway are intersected within and across species to find relations among the biological concepts. To identify candidate host molecular mechanisms associated with intestinal disease, we quantified overlap among sets of transcripts associated with mouse microbial abundance and human intestinal disease.

We determined that the abundance of *Roseburia* a bacterial genus from the *Lachnospiraceae* family (order *Clostridales*) is significantly correlated (p<0.05 and Spearman correlation ≥ 0.5) with the expression of *Cxcr4*, *Oma1*, *Igll1*, *Il1f8* in the CC-intestine (Table S3). Comparison of this set of correlated genes to those identified in published studies of intestinal disease transcriptomics in GeneWeaver revealed significant overlap with several ulcerative colitis and inflammatory bowel disease related gene sets (Jaccard Similarity p<0.000001) (Figure 3A). This analysis shows that *CXCR4: chemokine (C-X-C motif) receptor 4* is a biomarker of ulcerative colitis (UC) and Crohn’s Disease (CD) (Merelli et al., 2012; Simi et al., 1987) and the expression of the *Igll1* transcript can distinguish UC from CD in peripheral blood mononuclear cells of UC patients (Burczynski et al., 2006). Furthermore, this genetic analysis has now implicated two additional genes, *Il1f8* and *Oma1*, which are correlated with *Roseburia*, but not currently known to be associated with colitis. QTL *Micab4* regulated *Roseburia* a microbe that, at low intestinal abundance, is just one of the hallmarks of ulcerative colitis (UC) (Chen et al., 2014; Machiels et al., 2014; Willing et al., 2010).

The confidence interval for *Micab4* was 9.45 Mbp containing 95 genes (Fig 3B-C). At least five genes, *Id3* (probe18912 or ILMN_2687169), *Luzp1* (probe24933 or ILMN256903), *C1qa*, *C1qb*, and *C1qc* are compelling candidates for regulating *Roseburia* abundance because they either have cis-eQTL or they have an immune system function and contain variants with GWAS LOD scores >4 (Figure 3D). Therefore, we hypothesize that perturbation of a gene in the *Micab4* locus will impact *Roseburia* abundance through the indirect genetic regulation of *Cxcr4* and *Igll1* abundance. Of the five candidates, *Id3* and *C1qa* were testable in extant mouse models. *Luzp1* knock-out mice are perinatal lethal (Hsu et al., 2008). There was no evidence of intestinal pathology or abnormal abundance of *Roseburia* in *Id3* deficient mice (SUPP TABLE OF ABUNDACE IN ID3?). In contrast, comparison of wild-type mice to *C1qa* knockout mice, and mice with an intact but human derived *C1qa* gene reveals decreased abundance of *Roseburia* in the intestine of *C1qa* deletion mutants (Figure 4E). Mice with the intact human derived allele had an intermediate *Roseburia* abundance, and did not differ significantly from either the wild-type or C1qa deletion mutants. *C1qa*, *C1qb* and *C1qc* encode proteins that form the C1Q multimer in 1:1:1 ratio (Reid and Porter, 1976) and are reported to have synchronized transcription in some cells (Chen et al., 2011), therefore a polymorphism influencing one of them may influence the entire complex (Chick et al., 2016). SNP association mapping in the *C1qa* region reveals significant associations with variants for which B6, 129 and NZO alleles have one effect on *Rosburia* abundance and the other strains have an opposing effect. Several 3’UTR variants and synonymous variants have this segregation pattern. An additional SNP in *C1qa*, rs27625206 causes an amino acid change at AA16 changing a Thr (polar side chain) to an Ile (non-polar side chain) in A/J, NOD and PWK that may have functional effects. Collectively, these data suggest that genetic variation in the C1 complement system may influence the abundance of *Roseburia*, a colitis related bacterium.

**Figure 3.**
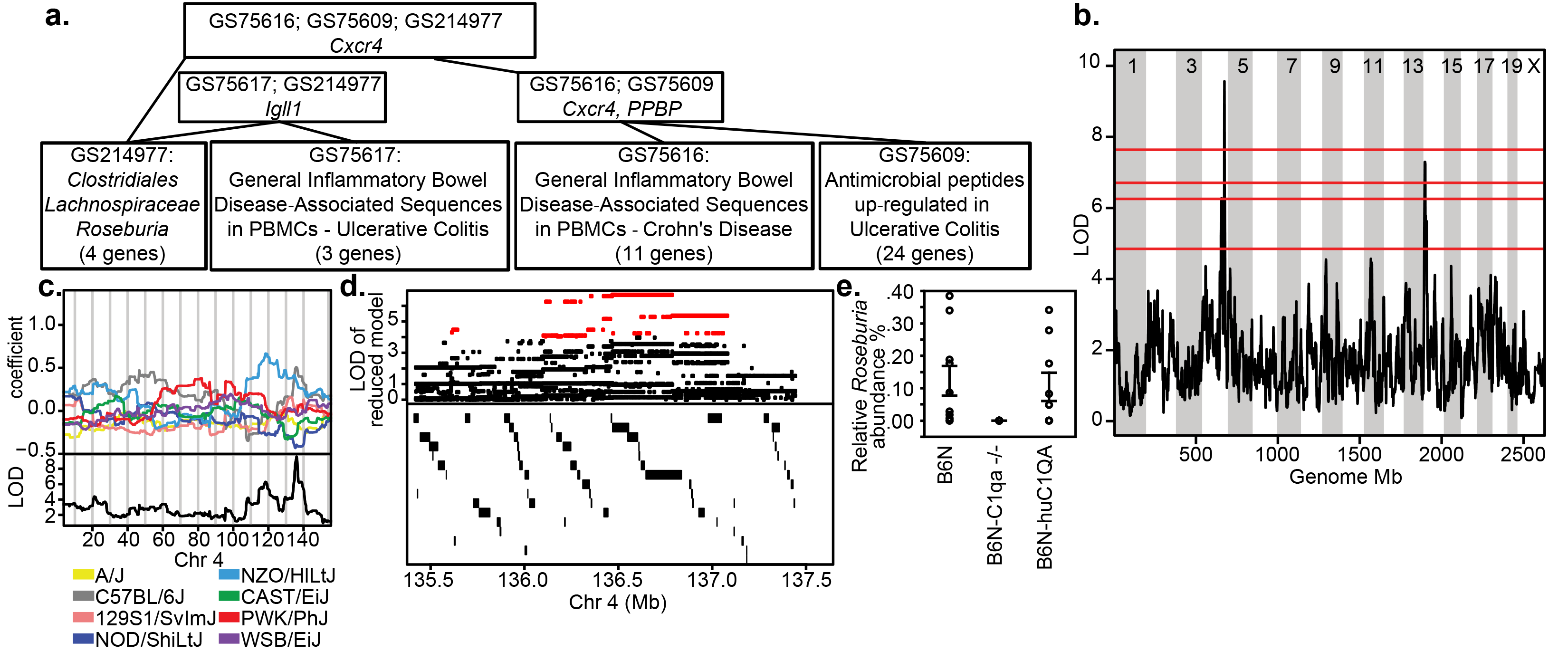
Hierarchical intersection of gene-microbe associations together with human gastrointestinal disorder literature. a) GeneWeaver output revealing hierarchical intersections of gene sets with the lowest level nodes representing individual gene-sets, and the higher level nodes representing 2-way, 3-way to n-way intersections of the inputs. The arrows show the direction of overlap of the genes. The genes associated with *Roseburia* (*Clostridiales Lachnospiraceae*) abundance include *Cxcr4*, *Oma1*, *Igll1*, *Il1f8*. *Cxcr4* has been implicated in two studies of UC and CD disease in humans and *Igll1* has been implicated in CD. b) Linkage mapping of microbial abundance using the additive haplotype model reveals a significant QTL (p<0.01) on chromosome 4. Horizontal lines represent permuted significance thresholds from the top down highly significant p<0.01, significant, p<0.05, highly suggestive, p<0.1, suggestive, p<0.63 c) Founder coefficients or allelic effects from the linkage model on chromosome 4 show the effects of each founder allele. d) The top panel shows the LOD score from association mapping in the QTL confidence interval. The bottom panel shows the genes and non-coding RNAs from the Mouse Genome Informatics database. e) Relative abundance of *Roseburia* in wild-type, C1qa deficient and humanized C1qa mice.

**Figure 4.**
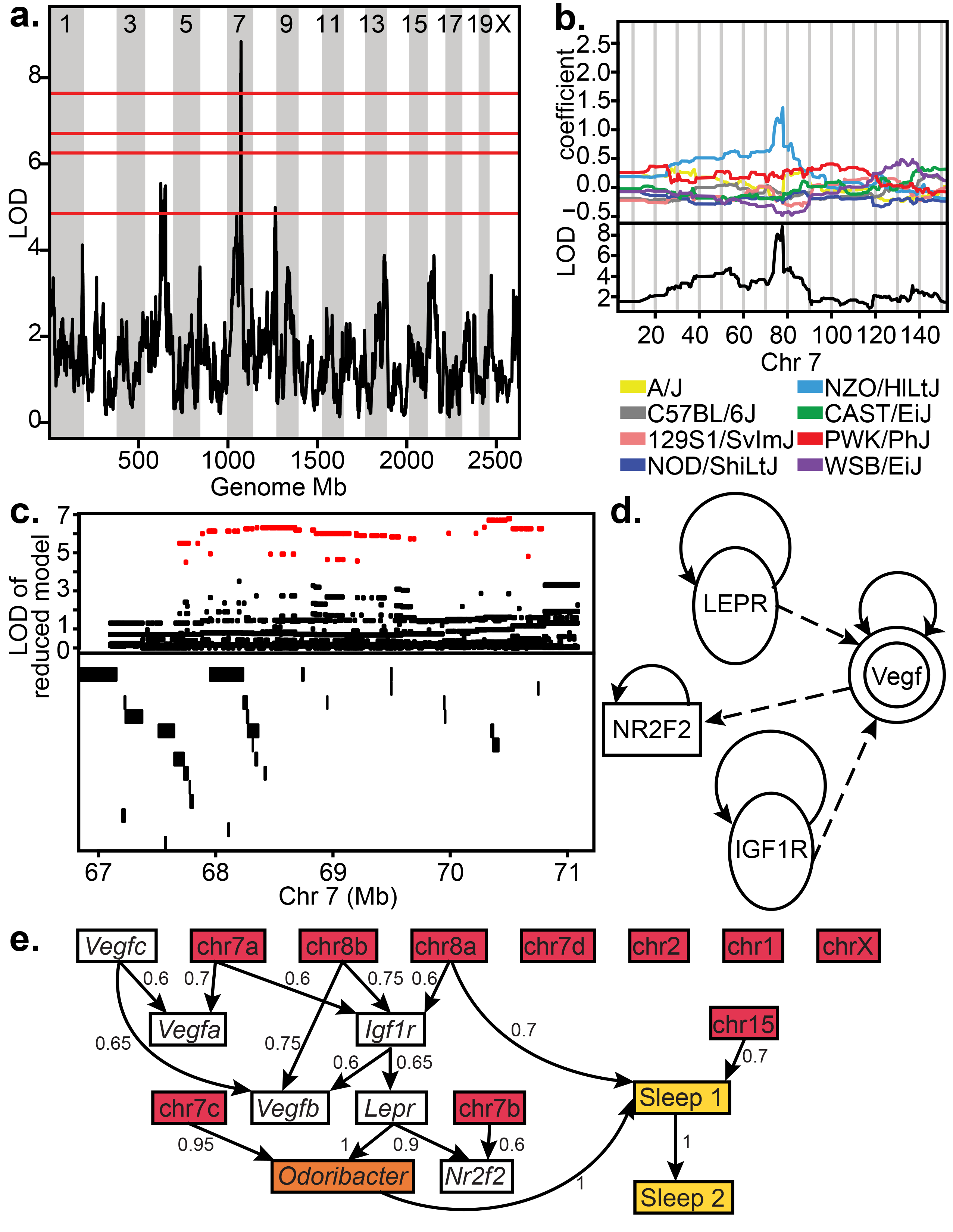
*Odoribacter* abundance in the cecum. a) Genome scan showing a significant QTL (p<0.01) peak on chromosome 7. Horizontal lines represent permuted significance thresholds from the top down highly significant p<0.01, significant, p<0.05, highly suggestive, p<0.1, suggestive, p<0.63 b) Detailed QTL map on Chromosome 7. Bottom: LOD score across Chromosome 7. Top: Allelic effect plots of eight coefficients of the QTL mixed model representing the effect of each CC founder haplotype on phenotype. The NZO allele on Chromosome 7 is associated with increased abundance of *Odoribacter* c) Top: LOD score for SNP association mapping in the QTL support interval (67.1-71.1). Red points indicate SNPs with significant association to *Odoribacter* abundance. Bottom: Genes and non-coding RNAs located in the QTL interval. d) Ingenuity Pathway Analysis of the positional candidate genes together with *Lepr* show a network path through *Vegf* and involving either *Nr2f2* or *Igf1r*. e) Inferred network relating sleep, microbe abundance, microbial abundance QTL, expression correlations and mutant mice. The network is a consensus representation of the 40 most likely BNs in a MCMC sample. Edge weights correspond to the marginal frequency of each directed edge in the top 40 Bayesian Networks. Sleep1 represents the sleep trait corresponding to the average of continuous sleep lengths over four full days, and sleep 2 represents the sleep trait Activity onset on the fourth day [h].

### Genetic correlation of microbial abundance to disease-related traits reveals a microbe associated with sleep

Correlation of disease-related traits with underlying biomolecular and microbial characteristics across individuals provides a powerful means to identify previously unknown mechanisms of disease. A total of 122 disease-related behavioral and physiological phenotypes were correlated using Kendall’s Tau with the abundance of each OTU, revealing 45 significant trait-microbe correlations (q< 0.05) (Storey, 2002) (Table 2). Of the trait-microbe pairs, 41 contained sleep phenotypes that showed significant associations with 10 different microbes. Among these, OTU 273 *Odoribacter* (ord. *Bacteroidales*, fam. *Porphyromonadaceae*) was the bacterium consistently correlated to the largest number of phenotypes (21), and also significantly correlated with decreased sleep time, among other sleep phenotypes (Table 2, Figure S4A).

**Table 2.**
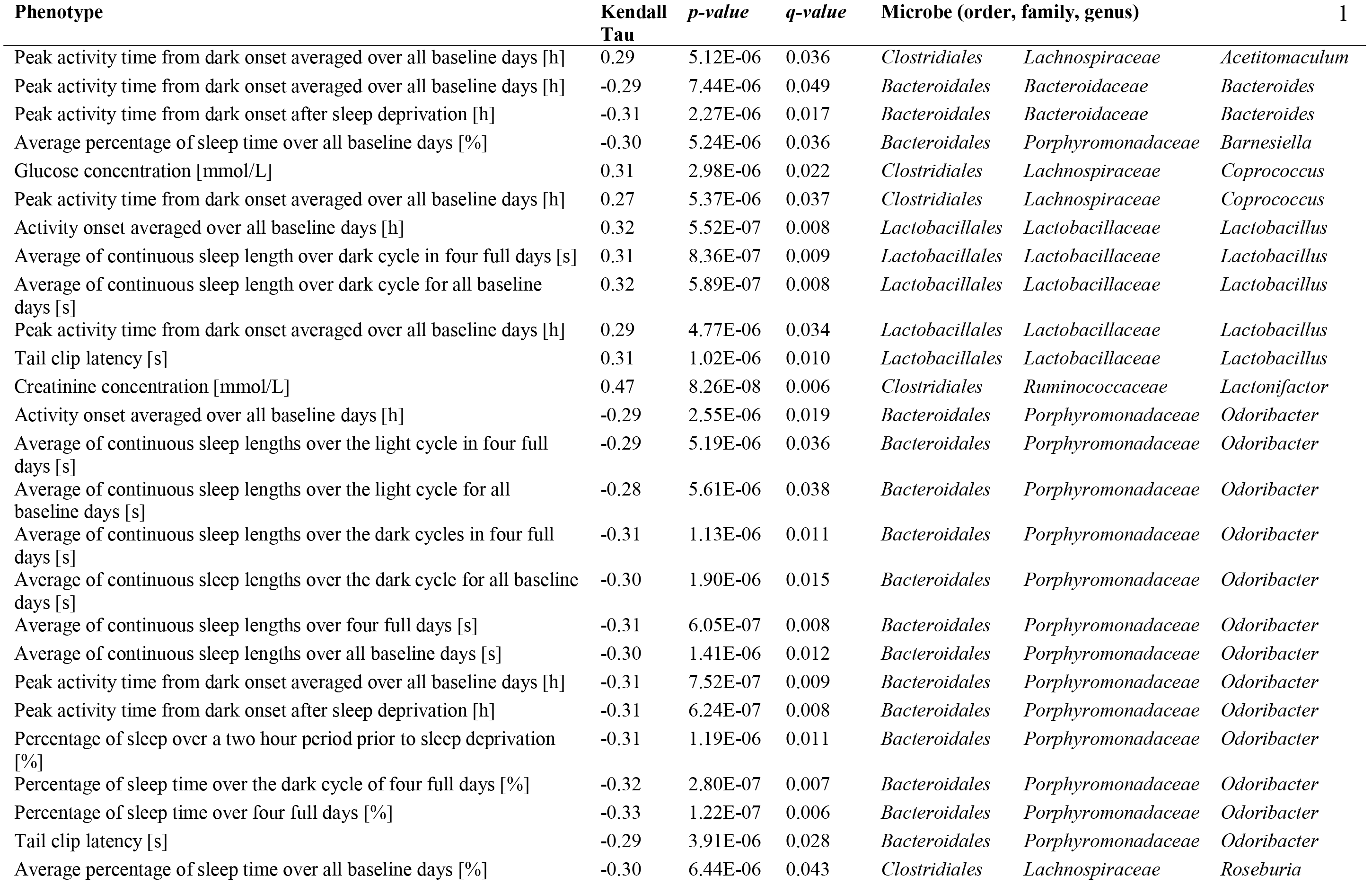
Correlations of disease related phenotypes to microbial abundance.

| Phenotype | Kendall<br>Tau | p-value | q-value | Microbe (order, family, genus) |  |  | 1 |
| --- | --- | --- | --- | --- | --- | --- | --- |
| Peak activity time from dark onset averaged over all baseline days [h] | 0.29 | 5.12E-06 | 0.036 | *Clostridiales* | *Lachnospiraceae* | *Acetitomaculum* |  |
| Peak activity time from dark onset averaged over all baseline days [h] | -0.29 | 7.44E-06 | 0.049 | *Bacteroidales* | *Bacteroidaceae* | *Bacteroides* |  |
| Peak activity time from dark onset after sleep deprivation [h] | -0.31 | 2.27E-06 | 0.017 | *Bacteroidales* | *Bacteroidaceae* | *Bacteroides* |  |
| Average percentage of sleep time over all baseline days [%] | -0.30 | 5.24E-06 | 0.036 | *Bacteroidales* | *Porphyromonadaceae* | *Barnesiella* |  |
| Glucose concentration [mmol/L] | 0.31 | 2.98E-06 | 0.022 | *Clostridiales* | *Lachnospiraceae* | *Coprococcus* |  |
| Peak activity time from dark onset averaged over all baseline days [h] | 0.27 | 5.37E-06 | 0.037 | *Clostridiales* | *Lachnospiraceae* | *Coprococcus* |  |
| Activity onset averaged over all baseline days [h] | 0.32 | 5.52E-07 | 0.008 | *Lactobacillales* | *Lactobacillaceae* | *Lactobacillus* |  |
| Average of continuous sleep length over dark cycle in four full days [s] | 0.31 | 8.36E-07 | 0.009 | *Lactobacillales* | *Lactobacillaceae* | *Lactobacillus* |  |
| Average of continuous sleep length over dark cycle for all baseline days [s] | 0.32 | 5.89E-07 | 0.008 | *Lactobacillales* | *Lactobacillaceae* | *Lactobacillus* |  |
| Peak activity time from dark onset averaged over all baseline days [h] | 0.29 | 4.77E-06 | 0.034 | *Lactobacillales* | *Lactobacillaceae* | *Lactobacillus* |  |
| Tail clip latency [s] | 0.31 | 1.02E-06 | 0.010 | *Lactobacillales* | *Lactobacillaceae* | *Lactobacillus* |  |
| Creatinine concentration [mmol/L] | 0.47 | 8.26E-08 | 0.006 | *Clostridiales* | *Ruminococcaceae* | *Lactonifactor* |  |
| Activity onset averaged over all baseline days [h] | -0.29 | 2.55E-06 | 0.019 | *Bacteroidales* | *Porphyromonadaceae* | *Odoribacter* |  |
| Average of continuous sleep lengths over the light cycle in four full days [s] | -0.29 | 5.19E-06 | 0.036 | *Bacteroidales* | *Porphyromonadaceae* | *Odoribacter* |  |
| Average of continuous sleep lengths over the light cycle for all baseline days [s] | -0.28 | 5.61E-06 | 0.038 | *Bacteroidales* | *Porphyromonadaceae* | *Odoribacter* |  |
| Average of continuous sleep lengths over the dark cycles in four full days [s] | -0.31 | 1.13E-06 | 0.011 | *Bacteroidales* | *Porphyromonadaceae* | *Odoribacter* |  |
| Average of continuous sleep lengths over the dark cycle for all baseline days [s] | -0.30 | 1.90E-06 | 0.015 | *Bacteroidales* | *Porphyromonadaceae* | *Odoribacter* |  |
| Average of continuous sleep lengths over four full days [s] | -0.31 | 6.05E-07 | 0.008 | *Bacteroidales* | *Porphyromonadaceae* | *Odoribacter* |  |
| Average of continuous sleep lengths over all baseline days [s] | -0.30 | 1.41E-06 | 0.012 | *Bacteroidales* | *Porphyromonadaceae* | *Odoribacter* |  |
| Peak activity time from dark onset averaged over all baseline days [h] | -0.31 | 7.52E-07 | 0.009 | *Bacteroidales* | *Porphyromonadaceae* | *Odoribacter* |  |
| Peak activity time from dark onset after sleep deprivation [h] | -0.31 | 6.24E-07 | 0.008 | *Bacteroidales* | *Porphyromonadaceae* | *Odoribacter* |  |
| Percentage of sleep over a two hour period prior to sleep deprivation [%] | -0.31 | 1.19E-06 | 0.011 | *Bacteroidales* | *Porphyromonadaceae* | *Odoribacter* |  |
| Percentage of sleep time over the dark cycle of four full days [%] | -0.32 | 2.80E-07 | 0.007 | *Bacteroidales* | *Porphyromonadaceae* | *Odoribacter* |  |
| Percentage of sleep time over four full days [%] | -0.33 | 1.22E-07 | 0.006 | *Bacteroidales* | *Porphyromonadaceae* | *Odoribacter* |  |
| Tail clip latency [s] | -0.29 | 3.91E-06 | 0.028 | *Bacteroidales* | *Porphyromonadaceae* | *Odoribacter* |  |
| Average percentage of sleep time over all baseline days [%] | -0.30 | 6.44E-06 | 0.043 | *Clostridiales* | *Lachnospiraceae* | *Roseburia* |  |

The *Micab7* on chromosome 7 regulates the relative abundance of *Odoribacter*. The QTL is 3.53 Mbp in size and contains 42 genes (Table 1). The allelic effects for each of the eight founder strain haplotypes are such that the NZO (New Zealand Obese) allele is associated with increased abundance (Figure 4A-C). This is significant because the NZO founder strain is obese and prone to a diabetes phenotype, and previous studies of gut microbiota in obesity and diabetes prone mice revealed that *Odoribacter*, *Prevotella and Rikenella* have been found in the microbiota of diabetic *db/db* (BKS.Cg-*Dock7*^m^ +/+ *Lepr*^db^/J) mice and are absent among db/+, +/+ littermates (Geurts et al., 2011). The *db/db* mice have also been shown to have abnormal sleep patterns in the form of altered sleep-wake regulation (Laposky et al., 2008).

We hypothesized that *Odoribacter*, *Lepr* and sleep are connected through a common mechanism. Specifically if the mechanism controlling altered sleep phenotype and the presence of *Odoribacter* in *Lepr* mutant *db/db* mice is the same mechanism that underlies the correlation of *Odoribacter* abundance and sleep in the CC mice, then we suspect overlap between one or more of the QTL positional candidates and the *Lepr* pathway, and that the perturbation of the gut microbiota of *db/db* mice should affect sleep patterns.

In order to investigate whether there is overlap between *Micab7* QTL positional candidate genes and *Lepr*, we performed an Ingenuity Pathway Analysis (IPA) of the 42 positional candidates, together with the gene *Lepr*. The most likely pathway from this database (Fisher’s Exact Test p < 10^−14^) contains the positional candidate genes *Nr2f2* and *Igf1R* interacting with *Lepr* through *Vegf* (Figure 4D).

Causal graphical models for phenotype-genotype networks (Rockman, 2008) were used to infer the direct and indirect associations among the results of the IPA, including *Lepr*, *Vegfa*, *Vegfb*, *Vegfc* the two positional candidates *Nr2f2* and *Igfl1r*, the leptin pathway and sleep. The network model also included abundance of *Odoribacter*, two sleep traits and the genotypes of the CC mice at the QTL. Bayesian Networks (BNs) are described by directed acyclic graphs (DAG), which can be efficiently decomposed and translated into the joint distribution of variables in the model (Koller and Friedman, 2009). Conditional Gaussian distributions were used to model the relationships between genotype and phenotype, and the network structure was learned using a Markov Chain Monte Carlo (MCMC) sampling scheme (Hageman et al., 2011), and averaging over the top structures (Hoeting et al., 1999). The graphical model is represented as a DAG which can be efficiently decomposed and translated into the joint distribution of variables in the model. If the QTL is associated with the regulation of the *Vegf* pathway, we would expect to see evidence for a network edge between the genotype and at least one of the two positional candidates, the downstream *Lepr* genes and the phenotype. Furthermore, this analysis can determine which positional candidate is most likely influenced by the causal variant. In aggregate summaries of the top 40 graphs, a repeatable relationship among the QTL, the positional candidate *Igf1r*, *Odoribacter* and sleep is observed (Figure 4E). This relationship is observed in the majority of graphs. Therefore there is a plausible interaction among the QTL, *Igf1r* abundance, the leptin pathway, *Odoribacter* and sleep.

### Broad spectrum antibiotic treatment alters sleep pattern in Lepr^db^ /Lepr^db^ mice

We then evaluated whether the presence of *Odoribacter* in *Lepr*^*db*^ mice could explain the altered sleep behavior reported in these mice. To eliminate *Odoribacter*, mice were given antibiotic treatment continuously from conception. As expected (Savage and Dubos, 1968) this broad spectrum treatment resulted in increased fecal contents of the cecum observed at dissection in both genotypes (Figure S3B), however, it also resulted in a genotype specific effect on sleep architecture. The percent sleep time for the antibiotic treated *db/db* mice over a 72 hour period showed a genotype × treatment interaction in a repeated measure MANOVA; Genotype × Treatment F_(71,45)_ = 2.1199, p = 0.0040 (Figure 5A-D). Fourier amplitude analysis of the cyclic activity between 4 and 7 hour periods (Figure 5E), showed a significant genotype × treatment effect (F_(3,120)_ = 12.2193, p < 0.0001) and individual LS Means Student t-test showed significant differences between control *db/db* and all three other groups (Table S4). V4 sequencing of cecal contents from *db/db* mice showed seven microbial taxonomic units that were absent in the antibiotic treated case and elevated in the water vehicle controls, including the two from the family containing *Odoribacter* (Figure 5F). Therefore, we conclude that *Odoribacter* abundance influences sleep architecture in a manner regulated by genetic variation in *Igf1r* through the *Vegf/Leptin* pathway.

**Figure 5.**
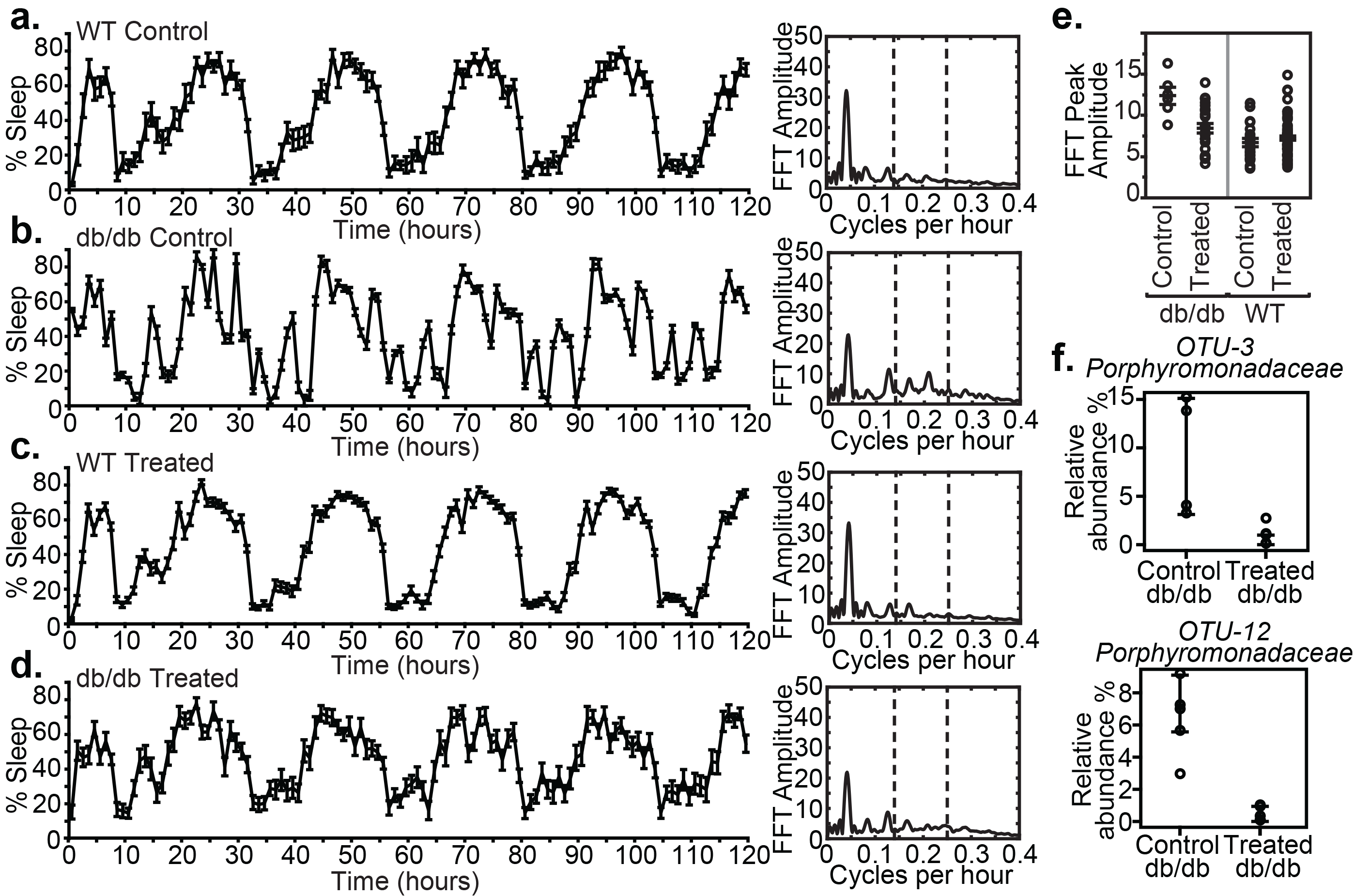
Mean and standard error for the percent time sleep over a 5 day test. a) Wild type (+/+ and db/+ genotypes) water only, b) Wildtype given antibiotics c) db/db mice water d) db/db mice given antibiotic Genotype and antibiotic treatment have a significant interaction affecting percent sleep time.e-h) FFT of sleep percentage time series. i) FFT Peak Amplitude corresponding to sleep percentage cycles with periods between 4 and 7 hours during the final 4 days of the sleep cycle, show significant genotype × treatment interaction as well as *db/db* control being significantly different from the other three groups. j-m) Microbes present in the control but absent in the antibiotic treated mice

## DISCUSSION

Using systems genetics and integrative functional genomics in the CC population, we traversed biological networks of genotypes, gene expression, microbes and disease-related phenotypes to identify host-microbial mechanisms underlying autism-, inflammatory bowel disease-and sleep-related phenotypes. The high allelic variation and precision of the incipient CC mouse population allowed us to map loci that control the abundance of 18 particular microbes, which could be further decomposed using SNP analysis, haplotype association and gene prioritization methods. Intestinal transcriptome profiling resulted in detection of ~1,600 significant eQTL and multiple clusters of transcripts and microbes whose abundances are jointly modified by genetic variation. Through genetic correlation network analysis, we relate these systems genetic networks to disease related phenotypes obtained in the same population of mice, and through cross-species integrative genomics, we relate the transcriptional correlates of microbial abundance to the transcriptional correlates of human disease.

Allelic variants influence the structure of microbial communities by creating conditions that promote or inhibit colonization by certain species (Spor et al., 2011). One way in which allelic variation manifests its effects is through the direct or indirect alteration of transcript abundance and the host environment, thereby impacting colonization. Other sources of variation may influence transcript abundance, including the presence of microbiota and their metabolites, disease states, and environmental variation. These sources of variation and their association with microbiota and disease can be detected through genetic correlation and probabilistic network analyses. By identifying network components and assessing causal relations among them through experimental perturbation, it is possible to understand the mechanisms of these relationships.

Using the incipient CC breeding population we have identified chromosomal loci across the genome, associated with the abundance of a variety of bacteria. We have identified eQTL in and clusters of microbes whose abundance correlates with the altered expression of sets of genes and disease related phenotypes. From each path of interrogation of the systems genetic network, we were able to confirm disease association and augment these observations with mechanistic insights into three disease areas. The systems genetic analysis of mouse intestinal microbiota enabled the discovery of multiple novel microbe-disease relationships. Some of these are supported by existing studies and others by new experimental evidence presented herein.

Previous work in humans has associated the specific microbial composition of the gut with autism-spectrum disorder, (De Angelis et al., 2013; Hsiao et al., 2013) and therefore, understanding how host and microbe interactions operate in a mouse model of ASD may lead to a better mechanistic understanding of the role of gut microbes in autism. The abundance of *Enterorhabdus* has been previously associated with valproic acid induced social behavior deficits and cecal short chain fatty acid metabolites (butyric acid) *(de Theije et al., 2014),* but the host mechanism mediating this effect and the presence of a naturally occurring host state that mimics the inducible model were unknown. Detecting a QTL for *Enterorohabdus* abundance and relating it to genetic regulation of *Trafd1* abundance enabled us to hypothesize and ultimately demonstrate that perturbation of *Trafd1* influences social behavior. *Trafd1* is known to influence inflammatory processes (Sanada et al., 2008) with broad effects on microbial colonization. Maternal infection during pregnancy and the resulting production of inflammatory molecules, has been shown to alter the fetal brain and produce autism-like behaviors in mice (Malkova et al., 2012). In the case of the VPA model of autism, *Trafd1* expression is increased by valproic acid (Jergil et al., 2011; Kultima et al., 2010), perhaps allowing for increased colonization by *Enterorhabdus* and resulting the differential microbial production of short chain fatty acids known to influence autism-related behavior (MacFabe, 2015).

Cross-species interrogation of intestinal disease genomics studies enabled extrapolation from mouse gene expression correlates of microbial abundance to differential gene expression in human gastrointestinal disorders. We identified gastrointestinal transcripts associated with the abundance of *Roseburia* (*Clostridiales, Lachnospiraceae)*, regulated by the *Micab4* QTL. The mouse locus containing the *Roseburia* abundance QTL on chromosome 4 is syntenic with a region of Chromosome 1 in human that has been repeatedly implicated in UC, IBD and Celiac disease (1p36.11 (Barrett et al., 2009; Dubois et al., 2010), 1p36.12 (Anderson et al., 2011; Barrett et al., 2009; Jostins et al., 2012) 1p36.13 (Anderson et al., 2011; Barrett et al., 2009; Ellinghaus et al., 2012; Franke et al., 2010; Jostins et al., 2012; Silverberg et al., 2009; Yang et al., 2013)). Therefore, finding the genetic variant underlying *Roseburia* abundance in the mouse may reveal mechanisms causing the dysbiosis seen in IBD. The locus contains complement proteins (*C1qa*, *C1qb*, *C1qc*), *Id3*, and *Luzp1*, compelling genetic candidates that also have high functional relevance due to their role in innate and adaptive immunity. Perturbation of *C1qa*, but not *Id3*, influences *Roseburia* abundance, and it is reasonable to speculate that other variants may exhibit similar effects.

Genetic correlation from mouse phenotype to microbial abundance enabled the identification of host and microbe influences on sleep architecture. The general role of microbes in sleep particularly in the cytokine response to infection is well documented (Krueger and Toth, 1994). Previous work in rabbits (Toth and Krueger, 1989) has shown that altered sleep patterns occur in response to an infectious challenge and that the sleep response is related to the type of infectious organism. Here we report for the first time the relationship between the abundance of a specific microbe and sleep. OTU273-*Odoribacter* (*Bacteroidales, Porphyromonadaceae*), regulated by the *Micab7* locus, was correlated with multiple sleep phenotype measures. Genomic network analyses revealed that the primary candidate gene for the QTL is *Igf1r*, a gene likely to function in the regulation of sleep as the somatotropic axis and IGF-1 signaling and sleep are intimately related (Obal et al., 2003). Perturbation of this pathway in the db/db *Lepr* mutant mouse is associated with elevated abundance of *Odoribacter* and an abnormal sleep phenotype, both of which we have shown can be restored to normal values through antibiotic treatment. The observation that indigenous microbes could affect sleep patterns suggests the potential for probiotic development in adjusting sleep patterns in those with clinical sleep disorders. Recent studies indicate a relationship between microbiota abundance and ultradian rhythms (Thaiss et al., 2014), and microbes of the *Odorbacter* were among five genera that decreased in the feces of intermittent hypoxia model of sleep apnea (Moreno-Indias et al., 2015).

In each of the above disease related studies, we utilized systems genetic networks to identify, model and validate the relationships among host genetics, genomics, microbiota and disease. The mouse provides an efficient, well-controlled system in which to employ this approach, though it is amenable to application in human populations. We demonstrated that using mouse genetics, we can identify relations that can be extrapolated to humans, though well-known issues in mechanistic conservation and direct translation must be considered. For example, despite high conservation across human and mouse genomes, specific biological mechanisms are not always entirely conserved, though the functional output of pathways and involvement in disease may be. Our approach to this challenge is to exploit network overlap, to identify elements of mouse networks that can be translated to human genetic and genomic networks, which we expect to function similarly but perhaps differ in the details of specific allelic variants, genetic mechanisms and particular microbiota involved. By developing our study around holistic quantitation of both host and microbe, in contrast to typical studies of individual perturbations and a specific focus on microbiota, we are able to generate broader networks amenable to integration and extrapolation to disease mechanisms. Much remains to be done in the functional validation of the conservation of these mechanisms.

In all studies of the interplay among host environment, microbiota and disease, the causal mechanisms underlying associations must be considered. Genetic variation influences the host environment creating conditions that are hospitable or inhospitable ecological niches for gut microbiota. Identifying the precise causal genetic variants underlying microbial composition is a lengthy process that has become more tractable with the advent of deep sequencing of the CC founders, high precision mapping populations including the Diversity Outbred derived from the CC, and the ability to integrate functional genomic data from other sources including epigenetic modification, non-coding variants, and disease associations.

By exploiting genetic heterogeneity among organisms, we were able to extract mechanistic relations among host, microbe and disease. The systems genetic strategy employed herein provides a wealth of data resources that can be further interrogated by investigators with an interest in specific host genes, variants, microbes and disease related phenotypes.

Furthermore, the strategy we present can be readily deployed in other genetically diverse populations to provide efficient, holistic assessment of microbial and host mechanisms of disease. Extracting these disease relevant mechanistic networks will provide insight into the complex interplay of host and microbe, revealing potential sources of disease etiology and points of therapeutic intervention.

## Acknowledgements

David Durtschi, Gene Barker and Ann Wymore performed genotyping and gene expression analysis for all samples. Darla Miller for her work with the Collaborative Cross at ORNL. Matthew Vincent produced the ‘deSNPed’ version of the Illumina probes for the CC founder strains. Jennifer Ryan, Neil Cole, Christine Rosales, Laura Anderson and Samantha Burrill conducted db sleep studies and dissections. Gaylynn Wells and Stacey Rizzo performed the 3-chamber social interaction test. Ashley Hunt and Amanda Ackovitz populated the GeneWeaver database with published studies of intestinal disease. This research was sponsored by the Laboratory Directed Research and Development Program of Oak Ridge National Laboratory, managed by UT-Battelle, LLC, for the U. S. Department of Energy under contract DE-AC05-00OR22725.

## Author Contributions

Jason A. Bubier wrote the paper and performed experiments, Vivek M. Philip designed the sample collection, genotyping and oversaw the analysis, Christopher Quince, James H. Campbell, Yanjiao Zhou, Tatiana Vishnivetskaya, Suman Duvvuru performed statistical analyses, Rachel Hageman-Blair performed probabilistic network modeling, Juliet Ndukum performed trait correlation analyses, Kevin D. Donohue and Bruce F. O’Hara designed the sleep system and associated algorithms, and assessed sleep data, Charles Phillips, Carmen M. Foster, Cymbeline T. Culiat oversaw genotyping analysis, David J Mellert assisted with data analysis and writing the manuscript, George Weinstock and Yanjiao Zhou performed the 16S sequencing and analysis in the follow up experiments, Erich J. Baker implemented the GeneWeaver system, Michael A. Langston developed graph algorithms and oversaw graph analysis, Anthony V. Palumbo, Mircea Podar supervised the gut pyrosequencing study and microbial clustering, and Elissa J. Chesler oversaw all aspects of the project and manuscript preparation.

## Supplementary material

**Figure S1 Cecal microbial profile across mouse samples.** Using 16S V4 rRNA gene sequence reads, a majority of cecum microbes belong to the Firmicutes phylum followed by Bacteroides. The other abundant phyla were Protobacteria (0.6%), Actinobacteria (0.6%).and Synergistetes (0.3%)

**Figure S2 eQTL map of cecum transcripts in CC mice.**

**Figure S3 Microbes differentially expressed in Trafd1 -/- mice compared to wild type controls.**

**Figure S4 Microbes and Sleep.** A) *Odiobacter* and average percentage of sleep time in dark cycle among CC lines. Kendal’s Tau = -0.3314, q-value=0.006104. B) Cecums of Control (left) and antibiotic treated (right) +/+ mice

**Table S1.** Founder strain intraclass correlations, sequences, and genus mapping based on Ribosomal Database Project for Operational Taxonomic Units found in at least 10% of CC mapping population

**Table S2.** Heritability and eQTLs for cecal intestinal transcript abundance in CC founders and mapping population for probes with strain ICC >.3

**Table S3.** GeneWeaver Gene Set IDs for sets of transcripts which are co-abundant with specific microbial taxa.

**Table S4.** Post-hoc comparison of mean sleep FFT peak amplitude in db/db and wild-type mice treated with broad spectrum antibiotic or vehicle control.

**Table S1.** Founder strain intraclass correlations, sequences, and genus mapping based on Ribosomal Database Project for Operational Taxonomic Units found in at least 10% of CC mapping population

**Table S2.** Heritability and eQTLs for cecal intestinal transcript abundance in CC founders and mapping population for probes with strain ICC >0.3

**Table S3.** GeneWeaver Gene Set IDs for sets of transcripts which are co-abundant with specific microbial taxa.

**Table S4.** Post-hoc comparison of mean sleep FFT peak amplitude in db/db and wild-type mice treated with broad spectrum antibiotic or vehicle control.

